# Targeted intragenic demethylation initiates chromatin rewiring for gene activation

**DOI:** 10.1101/2020.07.16.205922

**Authors:** Yanjing V. Liu, Mahmoud A. Bassal, Migara Kavishka Jayasinghe, Quy Xiao Xuan Lin, Chan-Shuo Wu, Jing Ping Tang, Junsu Kwon, Qiling Zhou, Hong Kee Tan, Alexander K. Ebralidze, Minh T.N. Le, Li Chai, Touati Benoukraf, Annalisa Di Ruscio, Daniel G. Tenen

## Abstract

Aberrant DNA methylation in the region surrounding the transcription start site is a hallmark of gene silencing in cancer. Currently approved demethylating agents lack specificity and exhibit high toxicity. Herein we show, using the *p16* gene as an example, that targeted demethylation of the promoter-exon 1-intron 1 (PrExI) region initiates an epigenetic wave of local chromatin remodeling and distal long-range interactions, culminating in gene-locus specific activation. Through development of CRISPR-DiR (DNMT1-interacting RNA), in which ad hoc edited guides block methyltransferase activity in a locus-specific fashion, we demonstrate that demethylation is coupled to epigenetic and topological changes. These results suggest the existence of a specialized “demethylation firing center (DFC)” which can be switched on by an adaptable and selective RNA-mediated approach for locus-specific transcriptional activation.

**One Sentence Summary:** Locus demethylation via CRISPR-DiR reshapes chromatin structure and specifically reactivates its cognate gene.

## Main Text

DNA methylation is a key epigenetic mechanism implicated in transcriptional regulation, normal cellular development, and function (*1*). The addition of methyl groups that occurs mostly within CpG dinucleotides is catalyzed by three major DNA methyltransferase (DNMT) family members: DNMT1, DNMT3a, and DNMT3b. Numerous studies have established a link between aberrant DNA methylation and gene silencing in diseases (*2, 3*).

Tumor suppressor gene (eg. *p16, p15, MLH1, DAPK1, CEBPA, CDH1, MGMT, BRCA1*) silencing is frequently associated with abnormal 5’CpG island (CGI) DNA methylation (*4*) and it is considered as the hallmark of most if not all cancers (*5*). Since 70% annotated gene promoters overlap with a CGI (*6*), the majority of studies have only concentrated on the correlation of CpG island promoter methylation and transcriptional repression, specifically focusing on the region just upstream of the transcription start site (TSS) (*2, 7–9*), but neglecting some studies showing the regulatory importance of regions downstream of TSS (*10, 11*). Therefore, the regulatory importance and mechanism of intragenic methylation on gene expression is still not clear.

Currently, demethylating agents are of limited utility, experimentally and therapeutically, because they act indiscriminately on the entire genome (*12*). Thus, an approach that is able to selectively modulate DNA methylation represents a powerful tool to study locus specific epigenetic regulation, and a potential non-toxic therapeutic option to restore expression of genes aberrantly silenced by DNA methylation in pathological conditions.

Epigenetic control hinges on a fine interplay between DNA methylation, histone modifications, nucleosome positioning, and their respective genetic counterparts: DNA, RNA, and distal regulatory sequences involved in the formation of specific topological domains (*13*). This structured organization is the driver for both gene activation or repression (*14*). It remains unclear to what extent locus-specific DNA demethylation contributes to chromatin structural rearrangements culminating in activation of silenced genes. To date, the lack of methods promoting naïve and localized specific demethylation have been the major constraint to understand the sequential mechanistic aspects enabling locus-specific activation.

Previously, our group (*15*) identified RNAs inhibiting DNMT1 enzymatic activity and protecting against gene silencing in a locus specific modality, termed DNMT1-interacting RNAs (DiRs). This interaction relies on the presence of RNA stem-loop-like structures, and is lost in their absence. By combining the demethylating features of DiRs with the targeting properties of the CRISPR-dead Cas9 (dCas9) system, we developed a novel platform, namely CRISPR-DiR, to induce precise locus specific demethylation and activation. The incorporation of DiR-baits into the single-guide RNA (sgRNA) scaffold enables the delivery of an RNA DNMT1- interacting domain to a selected location while recruiting dCas9 (*8, 16, 17*).

We selected *p16* as an example to test our platform, because it is one of the first tumor suppressor genes more frequently silenced by promoter methylation in cancer (*18*). We observed that gene-specific demethylation not only in the upstream promoter, but also in the exon 1- intron 1 region, is critical to initiate a stepwise process, followed by the acquisition of active chromatin marks (eg. H3K4Me3 and H3K27Ac) and decrease of silencing mark (H3K9me3), and interaction with distal regulatory elements, ultimately leading to stable gene-locus transcriptional activation. Overall, our studies point to discovery of a specialized promoter-exon 1-intron 1 region as a “demethylation firing center” responsive to RNA-mediated control and governing gene-locus transcriptional activation, elucidating the previously unknown importance of intragenic exon 1-intron 1 demethylation for active gene transcription.

## Results

### Development of the CRISPR-DiR system

The tumor suppressor gene *p16* (also known as *p16^INK4a^, CDKN2A*) is one of the first genes commonly silenced by aberrant DNA methylation in almost all cancer types, including hepatocellular carcinoma (HCC) (*4, 19, 20*), and therefore we chose it as a model to study the effect(s) of gene-specific demethylation. Studies on the secondary structure of the Cas9/dCas9- sgRNA-DNA complex revealed that the sgRNA scaffold can be modified without compromising the stability of the complex or its functionality. We hypothesized that incorporation of short loop sequences corresponding to L1 (Loop 1)and L2 (Loop 2) of the DiR we characterized before (*15*) could enable binding and inhibition of DNMT1 in a gene-specific manner (Fig. 1A). Several potential CRISPR-DiR designs were tested (Fig. 1B) to attain a platform in which 1) the sgRNA- dCas9 complex structure is stable; and 2) delivers efficient demethylation and gene activation.

**Fig. 1.**
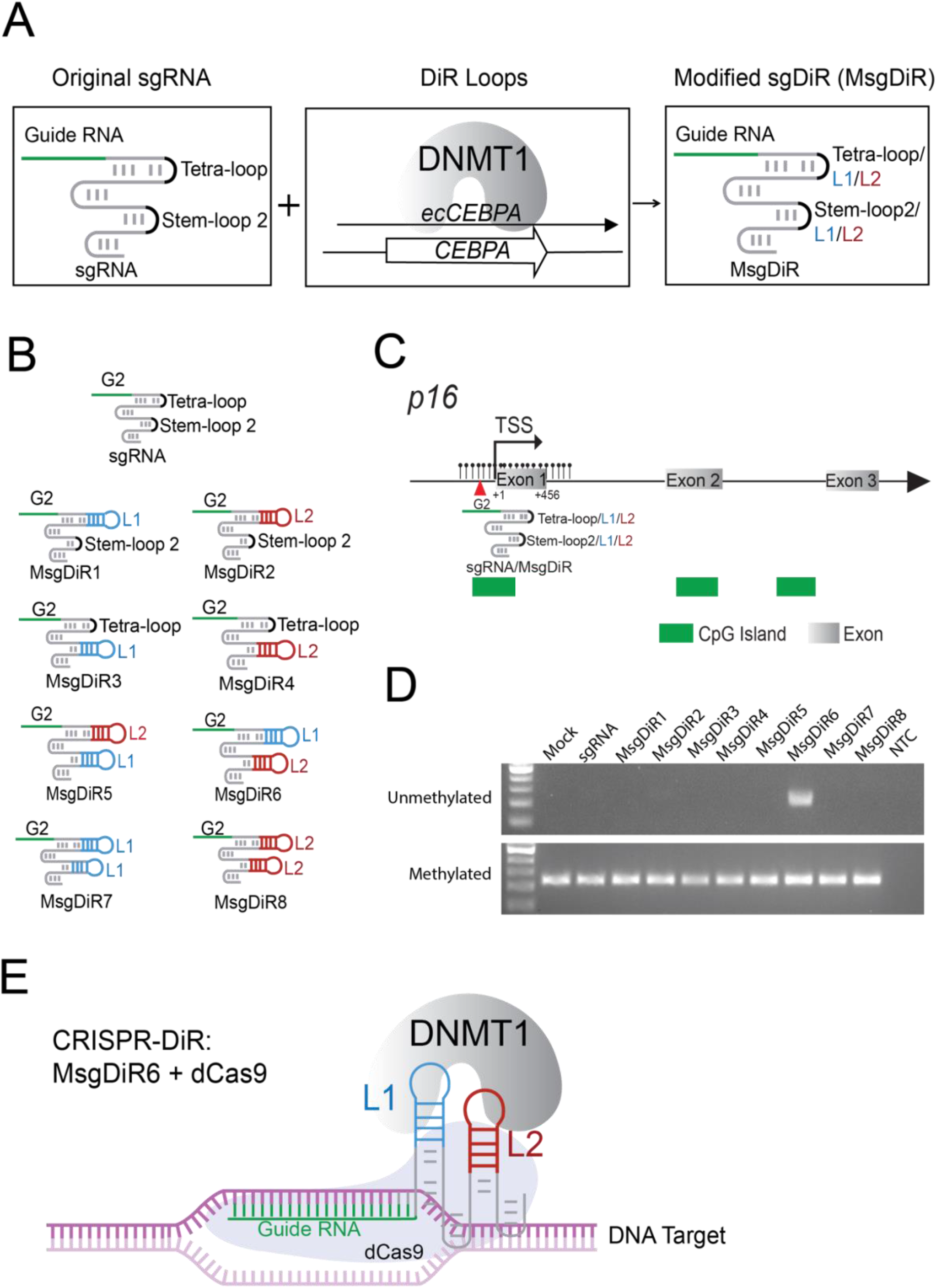
Development of the CRISPR-DiR system. (A) Rationale of CRISPR-DiR design. In the Modified sgDiR (MsgDiR), short DNMT1-interacting RNA (DiR) loops L1 and L2 from ecCEBPA were fused to the original sgRNA scaffold, tetra-loop and/or stem-loop 2 regions. (B) Diagrams of the original sgRNA control and eight different versions of MsgDiR design. All the sgRNA and MsgDiR constructs were utilized guide G2 targeting the *p16* gene proximal promoter. (C) Schematic representation of gene *p16* and the targeting site (G2) of sgRNA control and MsgDiRs. (D) Methylation Sensitive PCR (MSP) data demonstrating *p16* demethylation in SNU-398 cell lines 72 hours post-transfection. Mock: transfection reagents with H2O; sgRNA: co-transfection of dCas9+sgRNA (no DiR); Msg1-8: co-transfection of dCas9+MsgDiRs (with DiR) according to the design shown in Fig. 1C; NTC: none template control. (E) Schematic representation of the best CRISPR-DiR system after screening: dCas9+MsgDiR6, in which L1 is fused to sgRNA tetra-loop 2 while L2 is fused to sgRNA stem- loop 2.

Starting with a guide sequence (G2) successfully used in other studies, modified sgRNAs (MsgDiR) were designed in which the sgRNA scaffold embodied different combinations of L1 and L2 loops (Fig.1B) targeting the *p16* proximal promoter (Fig. 1C). dCas9 was co-transfected with either unmodified sgRNA or modified MsgDiR into SNU-398, a HCC cell line in which *p16* is methylated and silenced. Seventy-two hours after transfection, only the MsgDiR6 model induced *p16* demethylation (Fig. 1D). Further validation of MsgDiR6 with or without dCas9 in cells with either a non-targeting control guide (GN2) or *p16* guide (G2) for demethylation (Fig. S1A, S1B) demonstrated moderate activation of *p16* in dCas9 positive cells, while no effects were observed in absence of dCas9 (Fig. S1C). Intriguingly, MsgDiR6, which incorporates DiR loops L1 and L2 into the sgRNA scaffold (Fig. 1B, 1E, hereafter referred to as sgDiR), was the only design able to form a compatible predicted and functional secondary structure as reported for original sgRNA and other functional modified sgRNA design, suggesting that preserving the original structure is essential when editing the protruding loops in the sgRNA design.

In summary, we developed a novel functional platform, CRISPR-DiR, to induce locus-specific demethylation.

### CRISPR-DiR unmasks the *p16* transcriptional activator core

Although the initial analysis confirmed locus-specific demethylation, only a moderate activation of the mRNA was observed by the sgDiR (G2) targeting the *p16* proximal promoter upstream of transcription start site (TSS) (Fig. S1C). We sought to understand whether other than the promoter, the demethylation of additional intragenic regions within the locus were required for transcriptional activation. To identify demethylation-responsive elements, we decided to analyze the methylome of SNU-398 cells treated with the hypomethylating agent Decitabine (DAC), by Whole Genomic Bisulfite Sequencing (WGBS). We had expected demethylation in the well-studied upstream promoter region (Region D1). However, we detected a higher degree of demethylation within *p16* exon 1 (Region D2) and beginning of intron 1 (Region D3), suggesting a potential correlation between intragenic region demethylation and gene activation (Fig. 2A, 2B). To examine the contribution of the D2 and D3 regions on gene activation, we designed multiple sgDiRs specific to Region D1, D2, or D3, targeting either a single region individually or multiple regions in combination (Fig. 2C). sgDiRs targeting each region individually (Fig. 2C, S3A) could induce some degree of demethylation (Fig. S2B, S2C) and RNA production (Fig. 2D), with CRISPR-DiR targeting Region D2 leading to a greater than twofold increase in *p16* RNA (Fig. 2D).

**Fig. 2.**
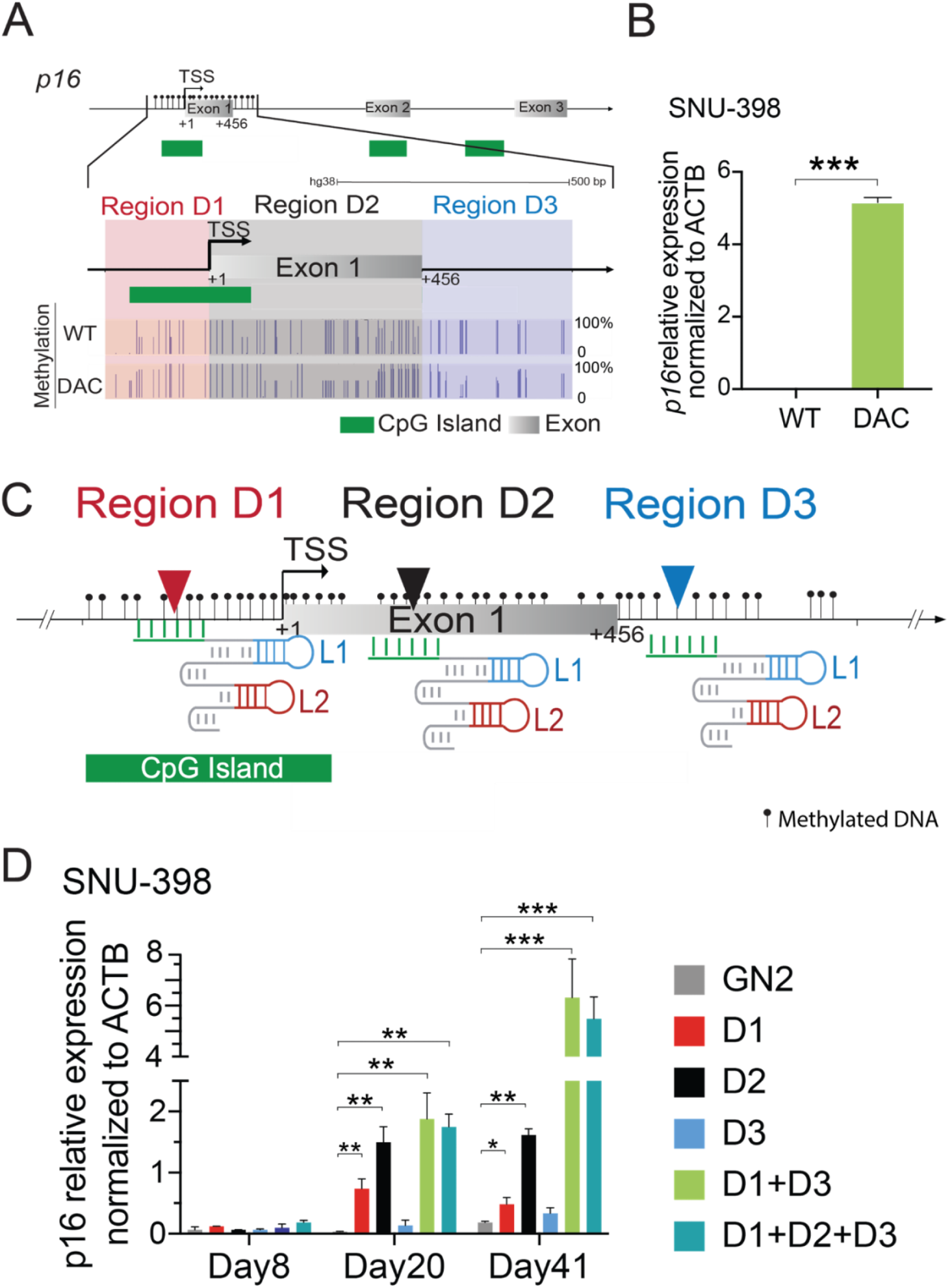
***p16* activation correlates with demethylation in exon 1 rather than promoter CpG island**. (A) Whole Genomic Bisulfite Sequencing (WGBS) results indicating the methylation profiles in the PrExI region (*p16* Promoter (Region D1)-Exon 1 (Region D2)-Intron 1 (Region D3) of wild type SNU-398 (WT) and SNU-398 treated with 2.5uM Decitabine for 72h (DAC). The height of the blue bar represents the methylation level of each CpG residue. (B) Real Time-Quantitative PCR (RT-qPCR) of *p16* gene expression in wild type and Decitabine treated SNU-398 cells, WT: wild type; DAC: Decitabine. (C) Schematic representation of the location of Region D1, Region D2 and Region D3 in the *p16* locus, as well as the CRISPR-DiR targeting sites in these three regions. (D) Real Time-Quantitative PCR (RT-qPCR) result of *p16* RNA in SNU-398 cell lines stably transduced with CRISPR-DiR lentivirus. Mean ± SD, n = 3, *P < 0.05; **P < 0.01; ***P < 0.001.

We further asked whether a) simultaneously targeting Region D1+D2+D3 or b) targeting demethylation in both the 5’ and 3’ ends (Region D1+D3) flanking a potential “seed” region D2, would lead to gene activation greater than any individual region alone. Indeed, the combined action of CRISPR-DiR targeting either Regions D1+D2+D3 or Region D1+D3 induced significantly greater increases in *p16* RNA than targeting any single region. Targeting Region D1+D3 achieved gene activation as great as targeting Region D1+D2+D3 all together (Fig. 2D), thus representing the simplest and most efficient targeting strategy for gene activation.

Collectively, these results demonstrate that the core epigenetic regulatory element of a gene is not necessarily contained within the promoter upstream of the TSS, but is augmented by the downstream exon 1 and adjacent intron 1 regions.

### CRISPR-DiR mediated intragenic demethylation is required for gene activation

The observation that the *p16* transcription pattern takes over a week to begin to change (Fig. 2D) in stably expressing CRISPR-DiR cells prompted us to trace the dynamic changes in *p16* demethylation over an extended period. Thus, *p16* demethylation and the respective gene expression was tracked for 53 days upon delivery of the most efficient targeting strategy, D1+D3, in SNU-398 cells. Bisulfite sequencing PCR (BSP) analyses revealed that demethylation initiated from regions D1 and D3 gradually increased from day 8 onwards, spreading to the intervening D2 region by Day 13 (Fig. 3A, 3B). Consistently, *p16* mRNA expression increased significantly after Day 13 (Fig. 3C), and p16 protein levels peaked after Day 20 (Fig. 3D), indicating that CRISPR-DiR initiated demethylation preceded transcriptional activation and protein expression. Strikingly, no demethylation “spreading” to surrounding regions (regions C and E) (Fig. S2A, S2D, S2E) was observed, suggesting that CRISPR-DiR mediated demethylation might be confined, and spreading exclusively within a regulatory core region (D2) (*21*). To demonstrate this effect was not confined to a single cell line, we delivered CRISPR-DiR into U2OS, a human osteosarcoma line with silenced *p16*, (Fig. 3E, 3F), and a similar trend in demethylation profiles and RNA expression was observed. In addition, no changes in RNA of the adjacent *p14* gene (located 20Kb upstream of *p16*, which is also methylated with no detectable expression), or *CEBPA* (located on another chromosome and actively expressed) was detected, thereby supporting the selectivity of the approach (Fig. S2F). In addition of the proof of principle in cell lines, we further performed xenograft study in mice. Using HCC cell line SNU- 398 as the model, 1) untreated, CRISPR-DiR 2) non-targeting (GN2) and 3) *p16* targeted cells were injected into mice and the tumor sizes were traced for 12 days (Fig. 3G). The result clearly indicated that the CRISPR-DiR approach can be applied in vivo, and the targeted activation of tumor suppressor gene *p16* effectively reduced tumor growth.

**Fig. 3.**
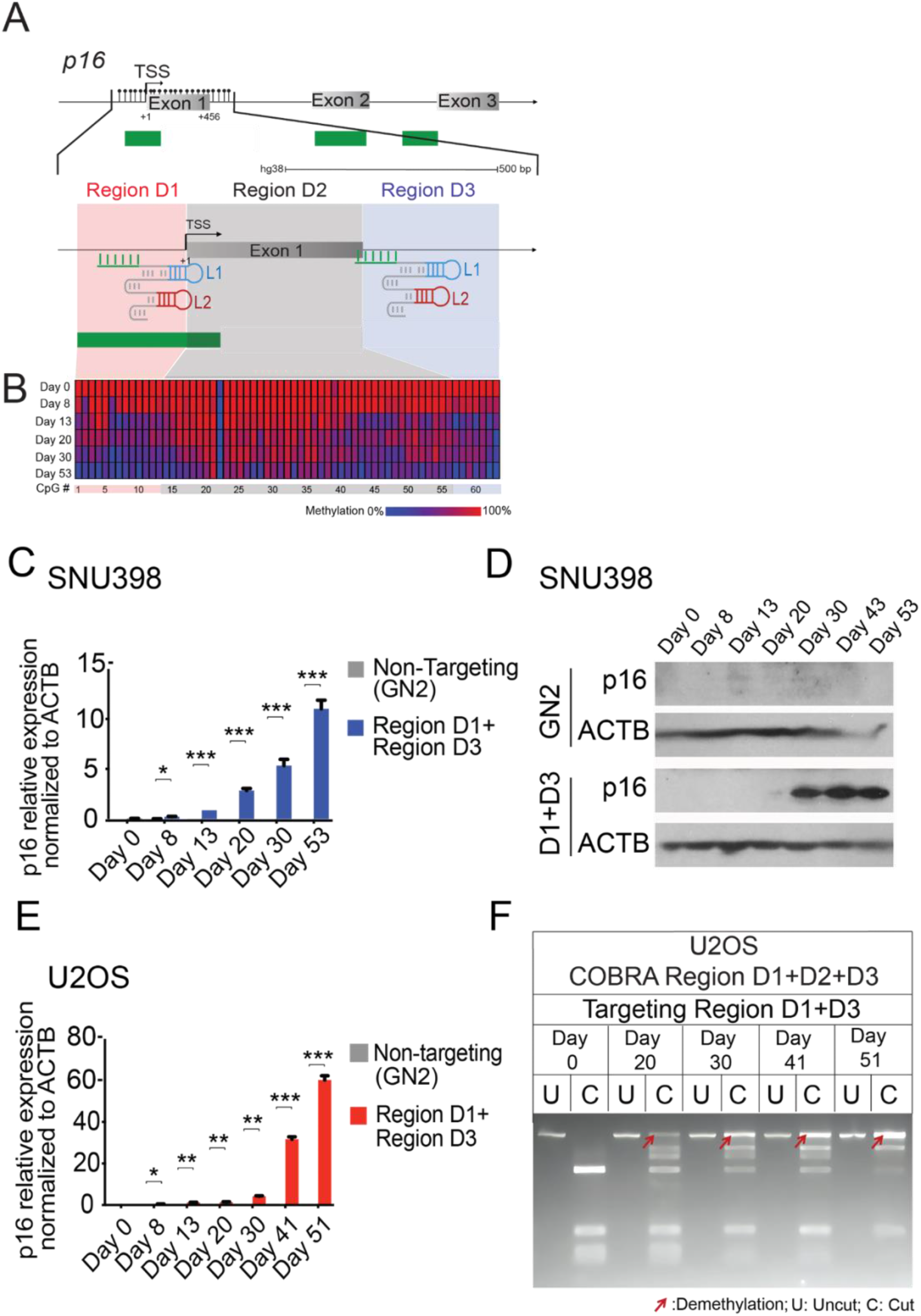
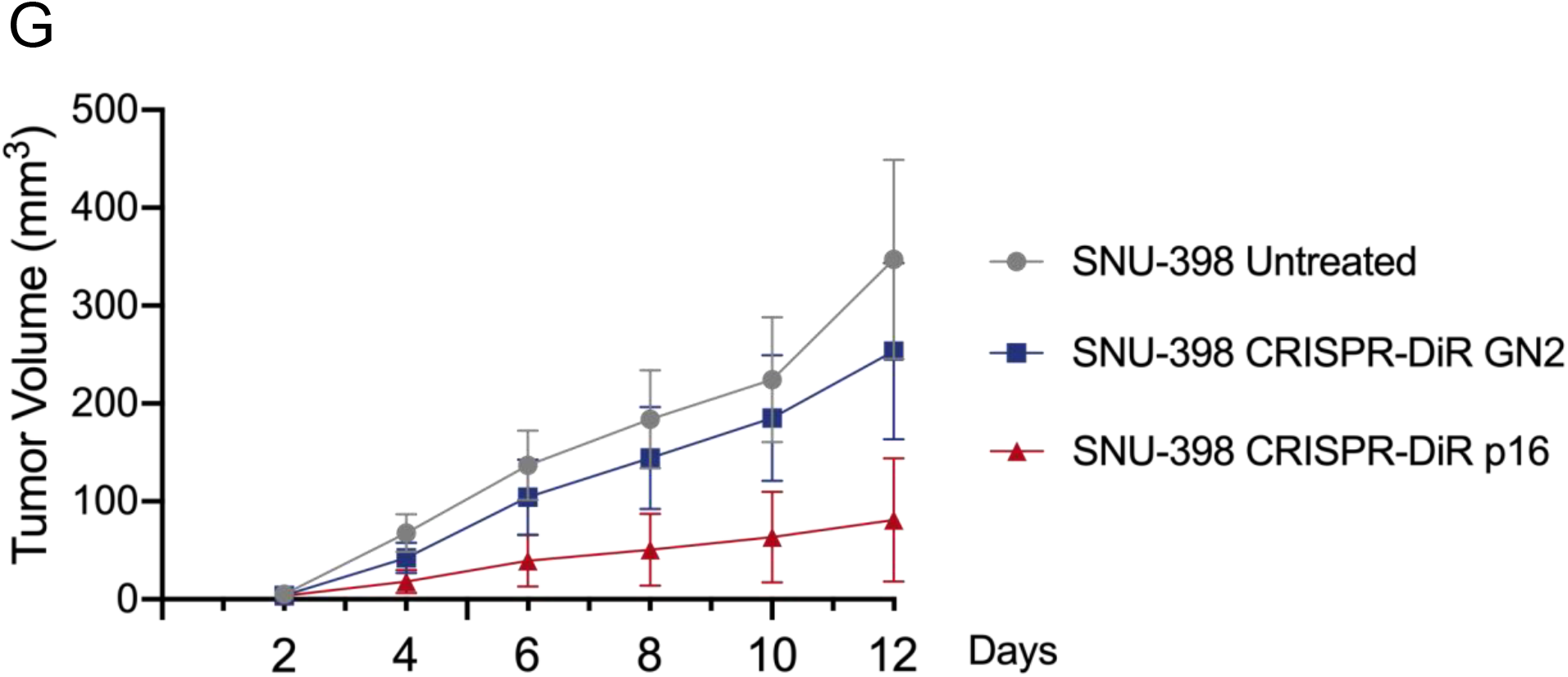
CRISPR-DiR targeting *p16* Region D1 and Region D3 simultaneously induced a dynamic process of demethylation and gene reactivation. (A) Schematic representation of the location of Region D1, D2, and D3 in *p16*, CRISPR-DiR targeting strategy: targeting *p16* Region D1 and Region D3 simultaneously. (B) Bisulfite Sequencing PCR (BSP) indicating the gradual demethylation profile in *p16* Region D1, D2, and D3 from Day 0 to Day 53 following CRISPR-DiR treatment in SNU-398 cells. (C) Real Time-Quantitative PCR (RT-qPCR) result showing *p16* mRNA expression after CRISPR-DiR treatment in SNU-398 cells. (D) Western Blot assessing p16 protein after CRISPR-DiR treatment. Beta actin (ACTB) was used as loading control. (E) RT-qPCR result showing *p16* gradual mRNA after the same CRISPR-DiR treatment in the human osteosarcoma U2OS cell line. (F) Combined Bisulfite Restriction Analysis (COBRA) representing the gradual demethylation profile in *p16* Region D1, D2, D3 (PrExI) from Day 0 to Day 53 with the same CRISPR-DiR treatment in U2OS cells. U=uncut, C= cut DNA. The band after cutting (lanes “C”) with migration equal to uncut represents demethylated DNA, and are indicated by red arrows. Mean ± SD, n =3, *P < 0.05; **P < 0.01; ***P < 0.001. (G) Inhibition of HCC tumor cell growth in vivo following CRISPR-DiR activation of *p16*.

### Targeted intragenic demethylation induces chromatin remodeling

To better evaluate whether demethylation by CRISPR-DiR, once initiated, was a lasting effect, we generated a Tet-On dCas9-stably expressing SNU-398 cell line, wherein dCas9 can be conditionally induced and expressed upon doxycycline addition. Within as soon as three days of induction treatment, *p16* demethylation and activation was observed and persisted for at least a month (Fig. 4A, 4B). These findings, along with our previous observations demonstrating steady increase in demethylation and RNA over nearly 2 months (Fig. 3B, 3C), pointed to the potential involvement of other epigenetic changes arising from the initial demethylation event and gene activation.

**Fig. 4.**
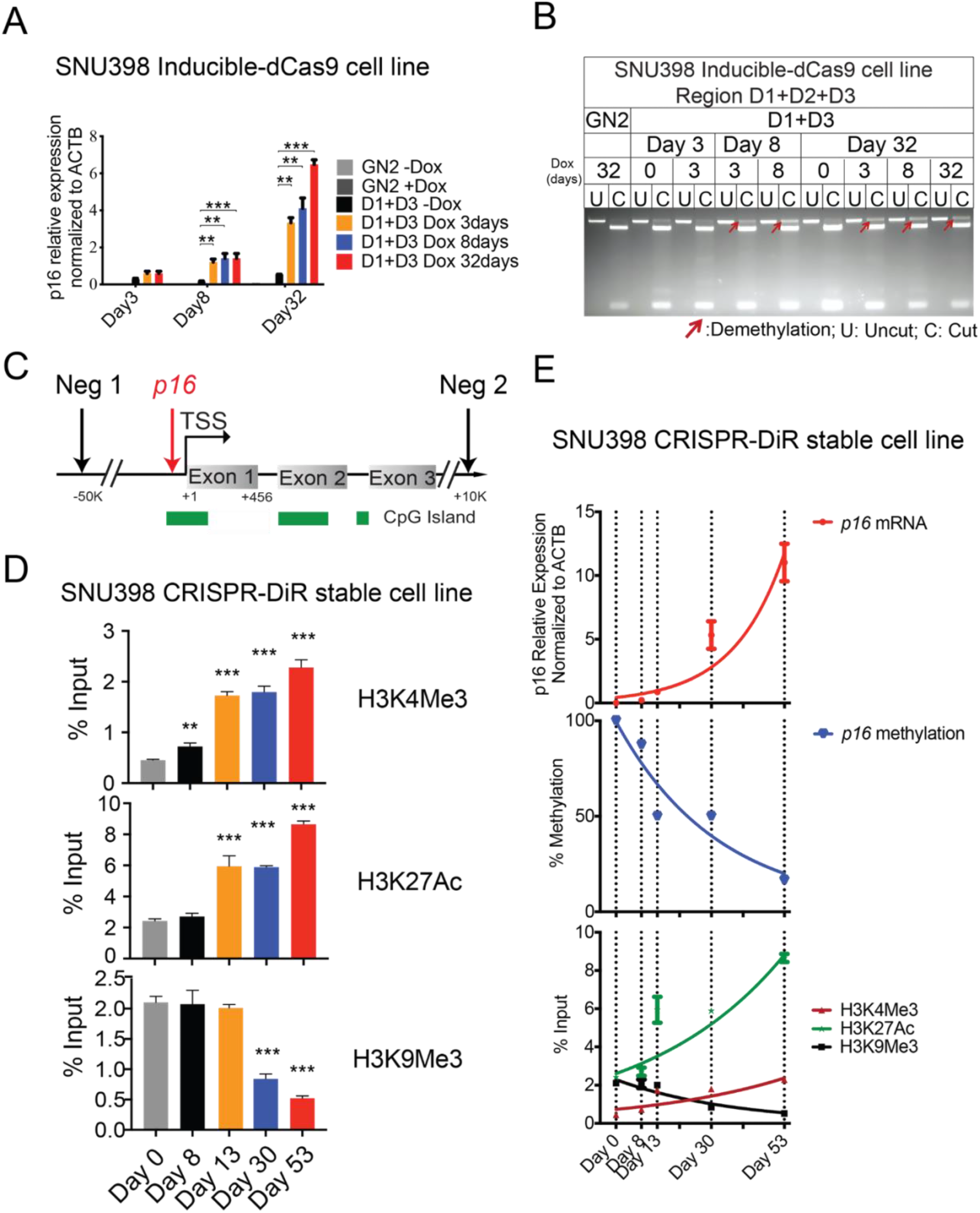
CRISPR-DiR effects are maintained for more than a month and PrExI demethylation leads to dynamic change in histone modifications. (A) Real Time-Quantitative PCR (RT-qPCR) result showing *p16* mRNA for more than a month in inducible CRISPR-DiR SNU-398 cells. In the inducible system, the same targeting strategy shown in Fig. 3A (Region D1 + Region D3) was used, and dCas9 expression was induced for 0 day, 3 days, 8 days, or 32 days following treatment with Deoxycytidine (Dox). All treatments were cultured and assayed at Day 0, Day 3, Day 8 or Day 32. (B) Combined Bisulfite Restriction Analysis (or COBRA) representing the demethylation profile of *p16* in inducible CRISPR-DiR SNU-398 cells. The demethylation status was maintained for more than a month with as short as three days induction. The band after cutting (lanes C) with equal migration as uncut (lanes U) represents demethylated DNA, indicated by red arrows. (C) Schematic representation of the location of ChIP-qPCR primers (Supplementary Table S7). Neg 1 and Neg 2: negative control primer 1 and 2 located 50 kb upstream and 10 kb downstream of *p16*, respectively. CpG island is indicated in green(D) ChIP-qPCR results showing the gradual increase in H3K4Me3 and H3K27Ac and decrease in H3K9Me3 enrichment in the *p16* PrExI region in SNU-398 cells stably transduced with CRISPR-DiR targeting D1 + D3 as in Fig. 3A. (E) Dynamic comparison of change in *p16* mRNA, methylation, and histone modifications in SNU-398 cells stably transduced with CRISPR-DiR targeting Region D1 + D3. Mean ± SD, n =3, *P < 0.05; **P < 0.01; ***P < 0.001.

We therefore hypothesized that loss of DNA methylation within the promoter-exon 1-intron 1 (PrExI) demethylation core region would facilitate histone changes and chromatin configuration for gene activation. To establish a direct correlation between these two events, Chromatin Immunoprecipitation (ChIP) with antibodies to the activation histone marks H3K4Me3 and H3K27Ac, or the repressive mark H3K9Me3, coupled with quantitative PCR (ChIP-qPCR) (Fig. 4C) was carried out in wild type and CRISPR-DiR treated (D1+D3) SNU-398 cells. We observed an enrichment of H3K4Me3 and H3K27Ac marks between Day 8 to 13 within the *p16* PrExI demethylation core region, inversely correlated with a progressive loss of the H3K9Me3 silencing mark (Fig. 4D, 4E), which corroborates the hypothesis that demethylation is the first event induced by CRISPR-DiR (Day 8), followed by gain of transcriptional activation marks in parallel to a loss of silencing marks (Day 8-13).

### Locally induced demethylation is essential to initiate distal long-range interactions

Thus far, we have shown that targeted demethylation of the p16 Mesa was sufficient to trigger potent gene expression. Additionally, our inducible CRISPR-DiR findings (**Figure 4a, 4b**) suggested that subsequent chromatin rewiring events might occur, resulting in lasting effects long after cessation of CRISPR-DiR induction.

A few studies have reported a *p16* enhancer region located ∼150 kilobases (kb) upstream of the *p16* TSS (*22–24*). Yet, long range interactions with the *p16* locus and the impact of locus-specific demethylation remains unexplored. To assess the impact of loss of DNA methylation on *p16* locus topology, we performed Circularized Chromosome Conformation Capture (4C) for CRISPR-DiR non-targeting (GN2) or targeting (D1+D3) Day 13 samples. We designed two viewpoints (‘baits’) as close to the promoter-exon 1-intron 1 demethylation core region as possible: Viewpoint 1, covering the exact targeted region D1 to D3 (Fig. 5A, 5B); while Viewpoint 2, covering the upstream promoter-exon 1 region (Fig. 5C, 5D). While Viewpoint 1 provides a closer examination of the targeted region, Viewpoint 2 overlaps more of the promoter. This two viewpoints design enables both an internal validation of the long-range interactions, and a careful analysis of the different interplay between distal regulatory elements and the promoter-exon 1 (viewpoint 2) or the exon 1-intron 1 (viewpoint 1) region, respectively (Fig. S3). Comparing the targeting (D1+D3) sample with the non-targeting (GN2) control, we indeed detected interaction changes between distal elements and the *p16* demethylated locus, scattered within 500 kb encompassing the *p16* targeted demethylation core region (PrExI) (Fig. 5B, 5D, S4). We further were able to identify the strongest interaction increases initiated by demethylation for both viewpoints (Fig. S3), which can represent potential distal enhancers for *p16* gene transcription. To note, among these strong interactions upon CRISPR-DiR induced demethylation, we not only detected novel interaction regions located more than 200kb upstream (E1), within the *Anril*- *p15-p14* locus (E3, E4), or more than 100kb downstream (E5, E6) of the *p16* TSS, but also observed interactions with the enhancer region previously described at ∼150kb upstream of *p16* TSS (E2) (*22–24*). These results indicated the reproducibility and reliability of the analysis, since the strong interactions overlap quite well between the two viewpoints (Fig. S3) and include the known enhancer regions. Furthermore, they point to potential novel *p16* enhancer elements and highlight a close interplay between *p16* and the neighboring gene loci *Anril-p15-p14*.

**Fig. 5.**
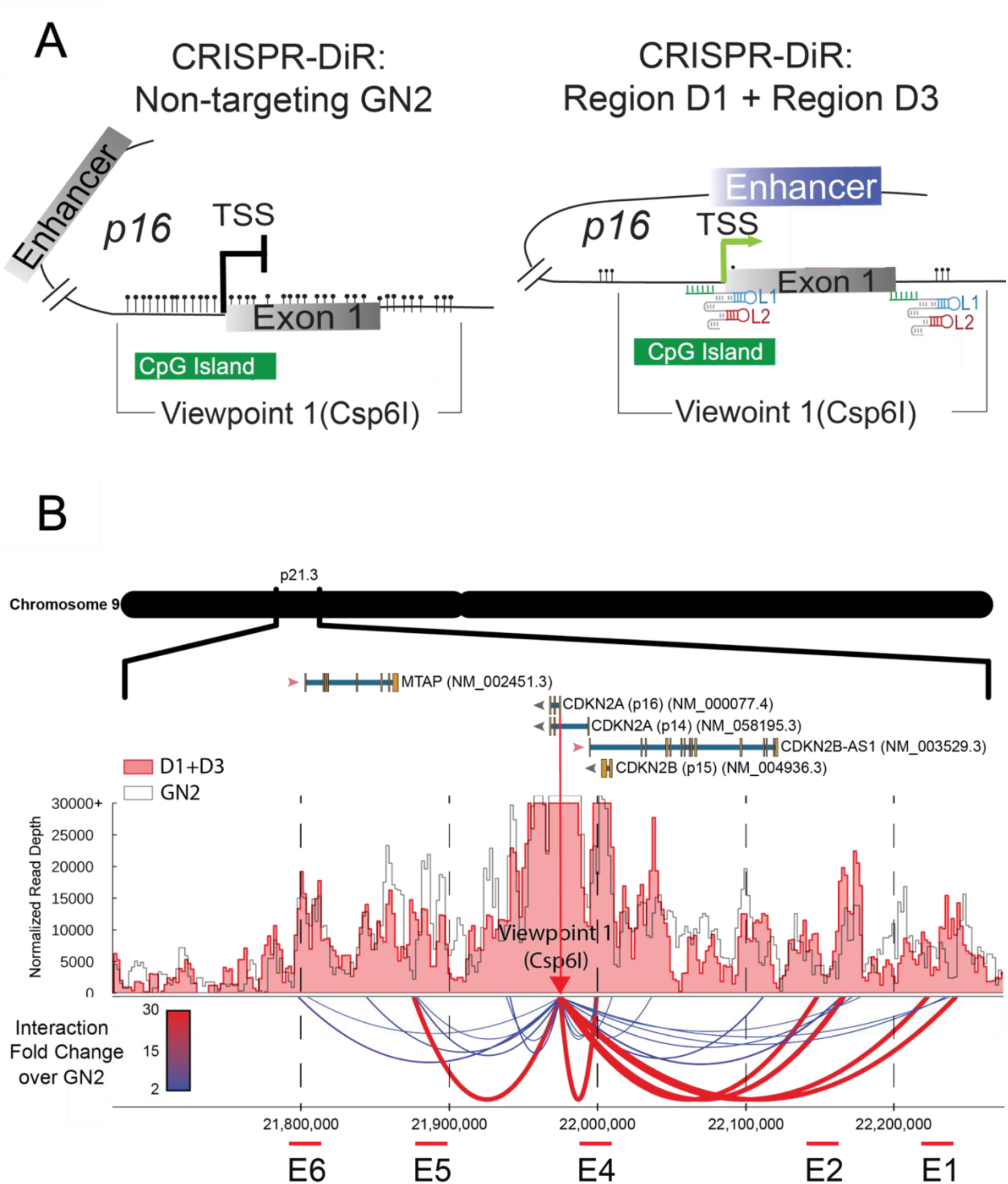
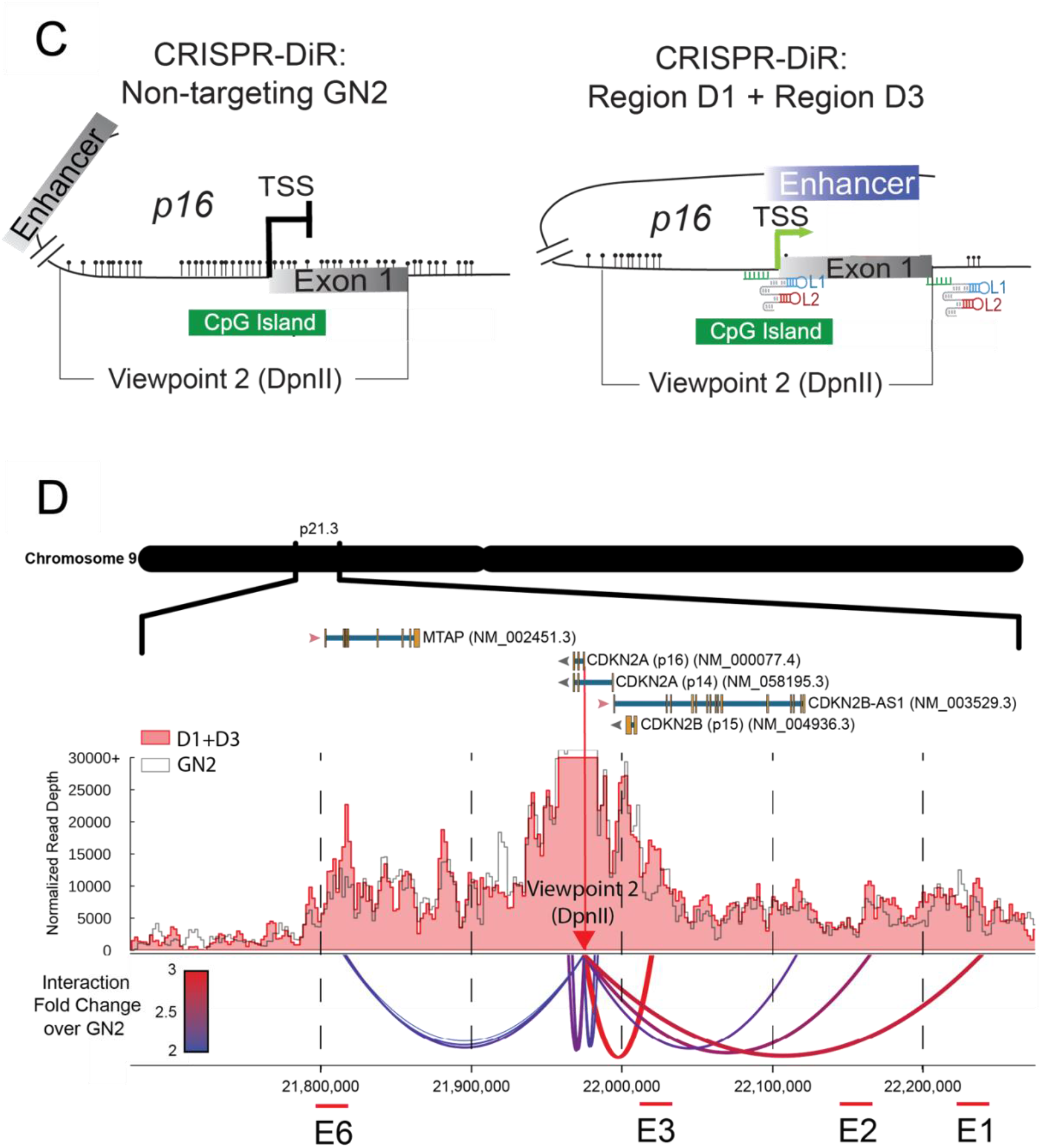
**CRISPR-DiR induced specific demethylation of *p16* PrExI remodels chromatin structure to activate gene expression**. (A) Hypothetical model of CRISPR-DiR induced demethylation of the PrExI region results in recruitment of distal regulatory elements, showing the 4C assay viewpoint 1 (generated by restriction enzyme Csp6I), covering the demethylated region (PrExI). (B) Circularized chromosome conformation capture (**4C**)-Seq analysis of CRISPR-DiR treated Day 13 samples (GN2 non-targeting control and D1+D3 targeted). Shown are interactions captured by 4C between *p16* viewpoint 1 and potential distal regulatory elements. The change in interaction was determined by normalizing the interactions of the targeted sample (D1+D3) to GN2 control; strong interaction changes are represented by the curves at the bottom from color blue to red, and the strongest interactions (potential distal enhancer elements) are highlighted and labelled as E1 to E6. (C, D) Same as panels A and B, except viewpoint 2 (generated by restriction enzyme DpnII), covering the 600 bp *p16* promoter region and *p16* exon 1, was used.

## Discussion

This study explores the functionalization of endogenous RNAs into an innovative locus-specific demethylation and activation technology named CRISPR-DiR (DNMT1-interacting RNA). By incorporating short functional DiR sequences into the scaffold of single guide RNAs, we generated a scalable, customizable, and precise system for naïve and localized demethylation and activation.

Using as a model the *p16* locus, a tumor suppressor gene frequently silenced by DNA methylation in cancers, we show that the core epigenetic regulatory element of gene activation is not contained within the extensively studied CpG-rich promoter region upstream of the *p16* TSS, but encompasses the proximal promoter-exon 1-intron 1 region (PrExI) (Region D1 to D3). By simultaneously engaging the 5’ (promoter) and 3’ (intron) regions flanking the activator core, our design introduces the requirements that enable consistent and effective demethylation spreading (Region D2, Fig. 3B) and that are missing in other CRISPR-based platforms targeting exclusively the immediate promoter (*7–9, 25*). Intriguingly, the CRISPR-DiR induced demethylation wave propagates inward into the middle of exon 1 region, while no demethylation is observed in the surrounding regions (Region C and E, Fig. S2D, S2E), despite the high CpG content, in contrast to what was previously suggested (*26, 27*). These findings demonstrate how demethylation of a critical regulatory core region is a necessary condition for gene activation for *p16*, which we have also demonstrated for the *SALL4* gene locus (*28*). The demethylation wave initiates a stepwise process followed by acquisition of active histone marks, decrease of silencing marks, and chromatin reconfiguration of the *p16* locus, ultimately steering long-range interactions with distal regulatory elements (Figs 5 and 6). In addition to a previously reported enhancer region located approximately 150 KB upstream of *p16* (*22–24*), we demonstrate that demethylation of the core region promotes interactions with a number of elements located as far as 500 kb away, indicating that localized and very specific adjustments of DNA methylation can broadly impact chromatin configuration and topological rearrangements (Fig. 6).

**Fig. 6.**
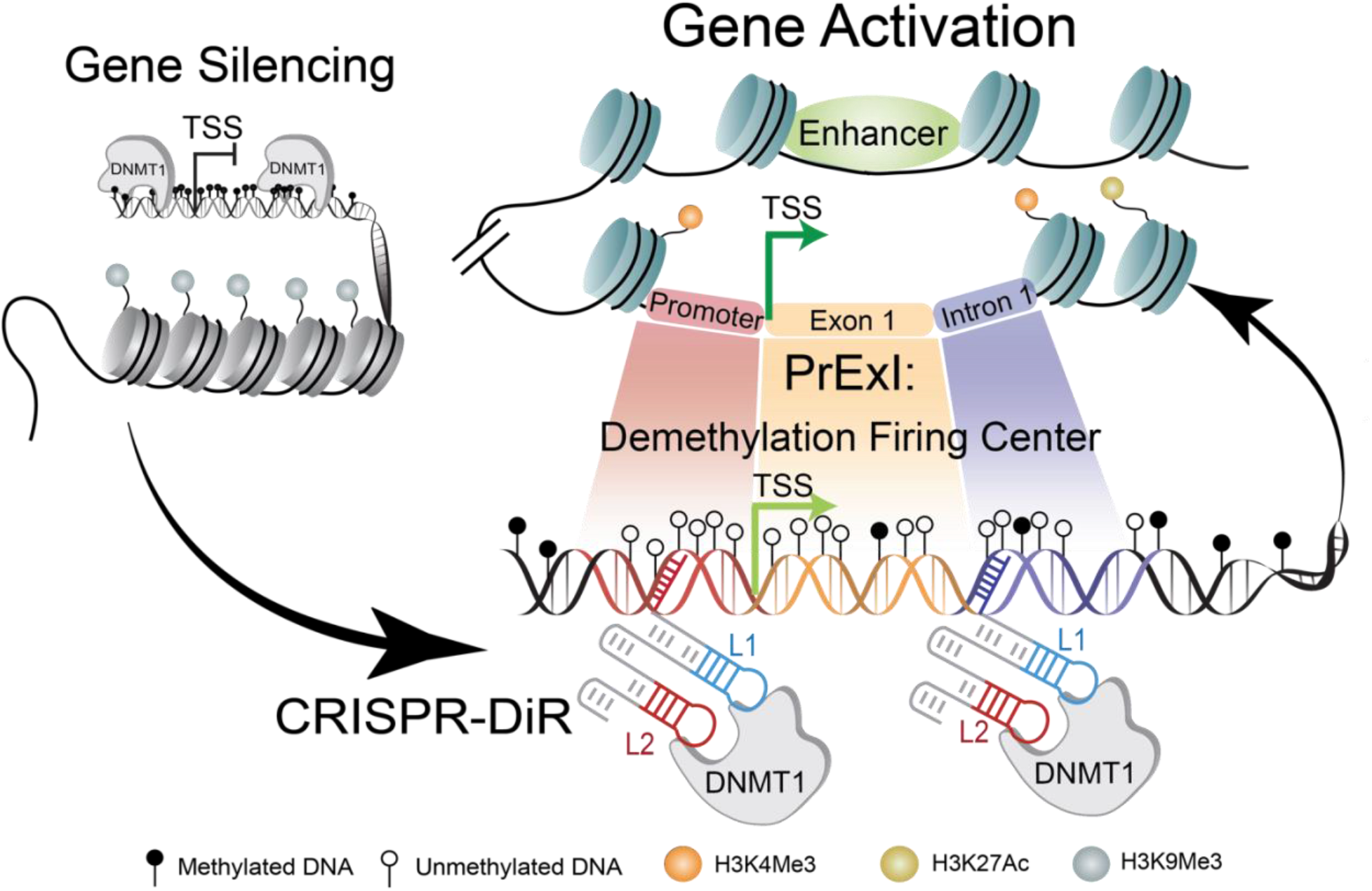
Schematic of CRISPR-DiR induced targeted demethylation in the Demethylation Firing Center (PrExI) initiating local and distal chromatin rewiring for gene activation. Gene silencing is coupled with aberrant DNA methylation in the region surrounding the transcription start site (TSS) as well as heterochromatin structure (upper left). Simultaneous targeting of the upstream promoter and beginning of intron 1 regions via CRISPR-DiR induces locus specific demethylation of the Demethylation Firing Center, which initiates an epigenetic wave of local chromatin remodeling and distal long-range interactions, culminating in gene-locus specific activation (on the right).

In conclusion, our data show how CRISPR-DiR induced demethylation of a small core element retained within 1 kb is able to propagate as far as 500 kb away, demonstrating the existence of an intragenic transcriptional initiator core which controls gene activation while acting as multiplier factor coordinating chromatin interactions. The features of this technology will aid in the identification of novel targets for clinical applications, developing alternative demethylation-based screening platforms, and designing therapeutic approaches to cancer or other diseases accompanied by DNA methylation.

## Materials and Methods

### Cell Culture

The human hepatocellar carcinoma (HCC) cell line SNU-398 was cultured in Roswell Park Memorial Institute 1640 medium (RPMI) (Life Technologies, Carlsbad, CA) supplemented with 10% fetal bovine serum (FBS) (Invitrogen) and 2 mM L-Glutamine (Invitrogen). Human HEK293T and human osteosarcoma cell line U2OS were maintained in Dulbecco’s Modified Eagle Medium (DMEM) supplemented with 10% fetal bovine serum (FBS). All cell lines were maintained at 37°C in a humidified atmosphere with 5% CO2 as recommended by ATCC and were cultured in the absence of antibiotics if not otherwise specified.

### RNA isolation

Total RNA was either extracted using AllPrep DNA/RNA Mini Kit (Qiagen, Valencia, CA) and treated with RNase-free DNase Set (Qiagen) following the manufacturer’s instructions, or isolated with TRIzol (Invitrogen). If the RNA isolation was carried out with the TRIzol method, all RNA samples used in this study were treated with recombinant RNase-free DNase I (Roche) (10 U of DNase I per 3 mg of total RNA; 37°C for one hour; in the presence of RNase inhibitor). After DNase I treatment, RNA samples were extracted with acidic phenol (pH 4.3) to eliminate any remaining traces of DNA.

### Genomic DNA extraction

Genomic DNA was extracted by either the AllPrep DNA/RNA Mini Kit (Qiagen, Valencia, CA) for BSP, MSP, and COBRA assays or by Phenol–chloroform method if extremely high-quality DNA samples were required for whole genomic bisulfite sequencing (WGBS). The Phenol–chloroform DNA extraction was performed as described (*29*). Briefly, the cell pellet was washed twice with cold PBS. 2 mL of gDNA lysis buffer (50mM Tris-HCl pH 8, 100mM NaCl, 25mM EDTA, and 1% SDS) was applied directly to the cells. The lysates were incubated at 65°C overnight with 2mg of proteinase K (Ambion). The lysate was diluted 2 times with TE buffer before adding 1mg of RNase A (PureLink) and followed by a 1 hour incubation at 37°C. The NaCl concentration was subsequently adjusted to 200mM followed by phenol-chloroform extraction at pH 8 and ethanol precipitation. The gDNA pellet was dissolved in TE pH 8 buffer.

### Quantitative Real-time PCR (qRT-PCR)

1 ug of RNA was reverse transcribed using Qscript cDNA Supermix (QuantaBio). The cDNAs were diluted 3 times for expression analysis. qRT-PCR on cDNA or ChIP-DNA were performed on 384 well plates on a QS5 system (Thermo Scientific) with GoTaq qPCR Master Mix (Promega, Madison, WI). The fold change or percentage input of the samples was calculated using the QuantStudio^TM^ Design & Analysis Software Version 1.2 (ThermoFisher Scientific) and represented as relative expression (ΔΔCt). All measurements were performed in triplicate. Primers used in this study are listed in Supplementary Table S1.

### 5-aza-2’-deoxycytidine (Decitabine) treatment

SNU-398 cells were treated with 2.5 μM of 5-aza-2’-deoxycytidine (Sigma-Aldrich) according to the manufacturer’s instructions. Medium and drug were refreshed every 24 h. RNA and genomic DNA were isolated after 3 days (72 h) treatment.

### Deoxycytidine (Dox) treatment

In the dCas9 Tet-On SNU-398 cells (inducible CRISPR-DiR system shown in Fig. 4A, 4B), the same targeting strategy shown in Fig. 3A (Region D1 + Region D3) was used, and dCas9 expression was induced following treatment with Deoxycytidine (Dox). Dox was freshly added to the culture medium (1 *µ*M) every day for Dox+ sample, while Dox- samples were cultured in normal medium without Dox. For Dox induced 3 days/8days samples, 1 *µ*M Dox was added to fresh medium for 3 days and 8 days accordingly, then the cells were kept in medium without Dox until Day 32; for Dox induced 32 days samples, 1 *µ*M Dox was added to fresh medium every day for 32 days. All treated cells were c*ul*tured and assayed at Day 3, Day 8 and Day 32.

### Transient transfections

SNU-398 cells were seeded at a density of 3.5 x 10^5^ cells/well in 6-well plates 24 hours before transfection employing jetPRIME transfection reagent (Polyplus Transfection) as described by the manufacturer. 2 μg mix of sgRNA/MsgDiR and dCas9 plasmid(s) (sgRNA/MsgDiR: dCas9 molar ratio 1:1) were transfected into each well of cells. The culture medium was changed 12 hours after transfection. Alternatively, the Neon™ Transfection System (Thermo Fisher) was used for cell electroporation according to the manufacturer’s instructions. The same plasmid amount and ratios were used in the Neon as in the jetPRIME transfection. The parameters we used for the highest SNU-398 transfection efficiency were 0.7 to 1.5 million cells in 100 µl reagent, voltage 1550 V, width 35 ms and 1 pulse. The culture medium was changed 24 hours after transfection. The plasmids used in this study are listed in Supplementary Table S2.

### Lentivirus production

pMD2.G, psPAX2, and lentivector (plv-dCas9-mCherry, pcw-dCas9-puro, plv-sgDiR-EGFP with different guide sequences) were transfected into 10 million 293T using TransIT-*LT1* reagent (Mirus), lentivector: psPAX2: pMD2.G 9 µg: 9 µg:1 µg. The medium was changed 18 hours post-transfection, and the virus supernatants were harvested at 48hr and 72hr after transfection. The collected virus was filtered through 0.45 µm microfilters and stored at -80 °C. The plasmids used in this study are listed in Supplementary Table S2. All the sgRNA, MsgDiRs scaffold sequences are listed in Supplementary Table S3, guide RNA sequences are listed in Supplementary Table S4 and the locations of each region (Region C, D1, D2, D3, and E) are listed in Supplementary Table S5.

### Generating CRISPR-DiR and inducible CRISPR-DiR stable cell lines

Both SNU-398 and U2OS cells were seeded in T75 flasks or 10cm plates 24 hours prior to transduction, and were first transduced with dCas9 or inducible dCas9 virus medium (thawed from -80 °C) together with 4 μg/mL polybrene (Santa Cruz) to make SNU398-dCas9, U2OS- dCas9, or inducible SNU398- dCas9 stable lines. Once incubated for 24 hours at 37°C in a humidified atmosphere of 5% CO2, the medium with virus can be changed to normal culture medium. The dCas9 positive cells were sorted using a mCherry filter setting with a FACS Aria machine (BD Biosciences) at the Cancer Science Institute of Singapore flow cytometry facility, while the inducible dCas9 positive cells were selected by adding puromycin at 2 µg/ml concentration in the culture medium every other day. The cells were further cultured for more than a week to obtain stable cell lines. Once the dCas9 and inducible dCas9 cell lines were generated, sgDiRs virus with different guide RNAs were mixed in equal volume and transduced into dCas9 or inducible dCas9 stable lines with the same method as described above. The sgDiRs used for generating each stable cell line, the location of each sgDiR, as well as the definition of Region D1, Region D2, and Region D3 can be found in Supplementary Table S4 and S5. All sgDiR stable cell lines were sorted using an EGFP filter with a FACS Aria machine (BD Biosciences) at the Cancer Science Institute of Singapore flow cytometry facility, and further assessed in culture by checking the efficiency by microscopy regularly.

### Western Blot Analysis

Total cell lysates were harvested in RIPA buffer (150mM NaCl, 1% Nonidet P-40, 50mM Tris, pH8.0, protease inhibitor cocktail) and protein concentrations were determined by Bradford protein assay (Bio-Rad Laboratories, Inc. Hercules, CA, USA) and absorbance was measured at 595 nm on the Tecan Infinite^®^ 2000 PRO plate reader (Tecan, Seestrasse, Switzerland). Equal amounts of proteins from each lysate were mixed with 3X loading dye and heated at 95 °C for 10 minutes. The samples were resolved by 12% SDS-PAGE (running buffer: 25 mM Tris, 192 mM Glycine, and 0.1% SDS) and then transferred to PVDF membranes (transfer buffer: 25 mM Tris, 192 mM Glycine, and 20% (v/v) methanol (Fischer Chemical)). Membranes were blocked with TBST buffer containing 5% skim milk one hour at room temperature with gentle shaking. The blocked membranes were further washed three times with TBST buffer and incubated at 4°C overnight with primary antibodies CDKN2A/p16INK4a (ab108349, Abcam, 1:1000), β-actin (Santa Cruz β-actin (C4) Mouse monoclonal IgG1 #sc-47778, 1:5000), followed by HRP-conjugated secondary antibody incubation at room temperature for one hour. Both the primary and secondary antibodies were diluted in 5% BSA-TBST buffer, and all the incubations were performed in a gentle shaking manner. The immune-reactive proteins were detected using the Luminata Crescendo Western HRP substrate (Millipore).

### Bisulfite treatment

The methylation profiles of the *p16* gene locus or the whole genome were assessed by bisulfite-conversion based assays. For DNA bisulfite conversion, 1.6-1.8 μg of genomic DNA of each sample was converted by the EpiTect Bisulfite Kit (Qiagen) following the manufacturer’s instructions.

### Methylation-Specific PCR (MSP), Combined Bisulfite Restriction Analysis (COBRA) and **bisulfite sequencing PCR (BSP)**

The bisulfite converted DNA samples were further analyzed by three different PCR based methods in different assays for the methylation profiles. For Methylation-Specific PCR (MSP), both methylation specific primers and unmethylation specific primers of *p16* were used for the PCR of the same bisulfite converted sample (the transient transfection samples in the sgRNA and MsgDiR1-8 screening assay). The PCR was performed with ZymoTaq PreMix (ZYMO RESEARCH) according to the manufacturer’s instructions, with the program: 95⁰C 10min, 35 cycles (95⁰C 30s, 56⁰C 30s, 72⁰C 1min), 72⁰C 7min, 4⁰C hold. Two PCR products of each sample (Methylated and Unmethylated) were obtained and analyzed in 1.5% agarose gels. For Combined Bisulfite Restriction Analysis (COBRA), primers specifically amplify both the methylated and unmethylated DNA (primers annealing to specific locus without any CG site) in each region were used for the PCR of the bisulfite converted samples. The PCR was performed with ZymoTaq PreMix (ZYMO RESEARCH) according to the manufacturer’s instructions, with the program: 2 cycles (95⁰C 10min,55⁰C 2min, 72⁰C 2min), 38 cycles (95⁰C 30s, 55⁰C 2min, 72⁰C 2min), 72⁰C 7min, 4⁰C hold. The PCR products were therefore loaded in a 1% agarose gel and the bands with predicted amplification size were cut out and gel purified. 400ng purified PCR fragments were incubated in a 20 µl volume for 2.5h-3h with 1 µl of the restriction enzymes summarized in Supplementary Table S6. 100ng of the same PCR fragments were incubated with only the restriction enzyme buffers under the same conditions as uncut control. The uncut and cut DNA were then separated on a 2.5% agarose gel and stained with ethidium the bromide. For bisulfite sequencing PCR, primers specifically amplify both the methylated and unmethylated DNA (primers annealing to the specific locus without any CG site) in Region D were used for the PCR of the bisulfite converted samples. The PCR was performed with ZymoTaq PreMix (ZYMO RESEARCH) according to the manufacturer’s instructions, with the program: 2 cycles (95⁰C 10min,55⁰C 2min, 72⁰C 2min), 38 cycles (95⁰C 30s, 55⁰C 2min, 72⁰C 2min), 72⁰C 7min, 4⁰C hold. PCR products were gel-purified (Qiagen) from the 1% TAE agarose gel and cloned into the pGEM-T Easy Vector System (Promega) for transformation. The cloned vectors were transformed into Stbl3 competent cells and miniprep was performed to extract plasmids for Sanger sequencing with either sequencing primer T7 or SP6. Sequencing results were analyzed using QUMA (Quantification tool for Methylation Analysis). Samples with conversion rate less than 95% and sequence identity less than 90% as well as clonal variants were excluded from our analysis. The minimum number of clones for each sequenced condition was 8. All the MSP, COBRA, and BSP primers as well as restriction enzymes can be found in Supplementary Table S6.

### Whole Genomic Bisulfite Sequencing (WGBS)

10 cm plates of wild type SNU-398 cells and Decitabine treated SNU-398 cells were washed twice with cold PBS. 2 mL of gDNA lysis buffer (50mM Tris-HCl pH 8, 100mM NaCl, 25mM EDTA, and 1% SDS) was added directly to the cells. The lysates were incubated at 65°C overnight with 2mg of proteinase K (Ambion). The lysate was diluted 2 times with TE buffer before adding 1mg of RNase A (PureLink) and followed by a one-hour incubation at 37°C. The NaCl concentration was subsequently adjusted to 200mM followed by phenol-chloroform extraction at pH 8 and ethanol precipitation. The gDNA pellet was dissolved in 1mL TE pH 8 buffer and incubated with RNase A with a concentration of 100ug/mL (Qiagen) for 1 hour at 37°C. The pure gDNA was recovered by phenol-chloroform pH 8 extraction and ethanol precipitation and dissolved in TE pH 8 buffer. 10ug of each gDNA samples (wild type and decitabine treated) were sent to BGI (Beijing Genomics Institute) for WGBS library construction and sequencing. The samples were sequenced to approximate 30X human genome coverage (∼90Gb) on a Hiseq X platform with 2X150 paired end reads.

### Xenograft in murine model

All experiments in mice were conducted according to our protocols approved by the Institutional Animal Care and Use Committee under National University of Singapore. Mice of similar ages were tagged and grouped randomly for control and test treatments. 2.5 million SNU- 398 HCC cells were injected into the flanks of NOD-SCID mice after transduction with 1) nothing (untreated), 2) a CRISPR-DiR system with non-targeting control guide (GN2), or 3) a CRISPR-DiR targeting region D1+D3 of *p16.* The tumor sizes were traced and recorded every two days for twelve days in total.

### Chromatin immunoprecipitation (ChIP)

ChIP was performed as described previously (*30*). Briefly, samples of 60 million cells were trypsinized by washing one time with room temperature PBS, then every 50-60 million cells were resuspended in 30ml room temperature PBS. Cells were fixed with 1% formaldehyde for 8 mins at room temperature with rotation. Excessive formaldehyde was quenched with 0.25M glycine. Fixed cells were washed twice with cold PBS supplemented with 1mM PMSF. After washing with PBS, cells were lysed with ChIP SDS lysis buffer (100 mM NaCl,50 mM Tris-Cl pH8.0, 5 mM EDTA, 0.5% SDS, 0.02% NaN3, and fresh protease inhibitor cOmplete tablet EDTA-free (5056489001, Roche) and then stored at -80 °C until further processing. Nuclei were collected by spinning down at 3000 rpm at 4°C for 10 mins. The nuclear pellet was resuspended in IP solution (2 volume ChIP SDS lysis buffer plus 1 volume ChIP triton dilution buffer (100 mM Tris-Cl pH8.6, 100 mM NaCl, 5 mM EDTA, 5% Triton X-100), and fresh proteinase inhibitor) with 10million cells/ml IP buffer concentration (for histone marker ChIP) for sonication using a Bioruptor (8-10 cycles, 30s on, 30s off, High power) to obtain 200 bp to 500 bp DNA fragments. After spinning down to remove debris, 1.2ml sonicated chromatin was pre-cleared by adding 50 µl washed dynabeads protein A (Thermo Scientific) and rotated at 4 ^0^C for 2 hrs. Pre-cleared chromatin was incubated with antibody pre-bound dynabeads protein A (Thermo Scientific) overnight at 4 ^0^C. For histone marker antibodies, 50 μl of Dynabeads protein A was loaded with 3 μg antibody. The next day, magnetic beads were washed through the following steps: buffer 1 (150 mM NaCl, 50 mM Tris-Cl, 1 mM EDTA, 5% sucrose, 0.02% NaN3, 1% Triton X-100, 0.2% SDS, pH 8.0) two times; buffer 2 (0.1% deoxycholic acid, 1 mM EDTA, 50 mM HEPES, 500 mM NaCl, 1% Triton X-100, 0.02% NaCl, pH 8.0) two times; buffer 3 (0.5% deoxycholic acid, 1 mM EDTA, 250 mM LiCl, 0.5% NP40, 0.02% NaN3) two times; TE buffer one time. To reverse crosslinks, samples were incubated with 20μg/ml proteinase K (Ambion) at 65°C overnight. The samples were then extracted with phenol:chloroform:isoamyl alcohol (25:24:1) followed by chloroform, ethanol precipitated in the presence of glycogen, and re-suspended in 10mM Tris buffer (pH 8). After reverse crosslinking and purification of DNA, qPCR was performed with the primers listed in Supplementary Table S7. Briefly, *p16* primer detecting the enrichments of all histone markers is located in the proximal promoter region within 100 bp around TSS; primers located 50 kb upstream of *p16* (Neg 1) and 10 kb downstream of *p16* (Neg 2) are the negative control primers located in the regions without enrichment of any of the above proteins. The antibodies used in ChIP assays were: H3K4Me3 (C42D8, #9751, Cell Signaling Technologies), H3K27Ac (ab45173, Abcam), H3K9Me3 (D4W1U, #13969, Cell Signaling Technologies), - Rabbit IgG monoclonal (ab172730, Santa Cruz).

### Circularized Chromosome Conformation Capture (4C)- Seq

4C-seq was performed as described previously (*31*) with modifications (*32*). In brief, SNU398 cells with stable CRISPR-DiR treatment for 13 days were used for 4C-Seq. 30 million sample a) guided by GN2 non-targeting and 30 million sample b) guided by guides targeting region D1+D3 were crosslinked in 1% formaldehyde for 10 mins at RT with rotation. Then formaldehyde was neutralized by adding 2.5 M glycine to a final concentration of 0.25 M and rotating for 5 mins at RT. After washing in cold PBS, cells were resuspended in 9 ml lysis buffer (10mM Tris-HCl pH8.0, 10mM NaCl, 5mM EDTA, 0.5% NP 40, with addition of EDTA-free protease inhibitor (cOmplete tablet, freshly dissolved in nuclease free water to make a 100X stock, 5056489001, Roche) and lysed multiple times with resuspension every 2-3 mins during the 10mins incubation on ice. After lysis, each lysate was split into two 15ml falcon tubes for viewpoint 1 (Csp6I) or viewpoint 2 (DpnII), respectively (4.5ml/tube, 15 million cells). After spinning down at 3,000 rpm for 10 mins at 4 ^0^C, each nuclear preparation was washed with 500 µl 1X CutSmart buffer from NEB and spun at 800g for 10 min at 4^0^C, followed by resuspension into 450 µl nuclease free (NF) H2O and transferring exactly 450 µl of the sample into a 1.5mL eppendorf tube. To each tube, 60 µl of 10X restriction enzyme buffer provided together with the corresponding restriction enzyme (viewpoint 1: 10X Buffer B (ER0211, Invitrogen); viewpoint 2: 10X NEBuffer™ DpnII (R0543M, NEB)) and 15 µl of 10% SDS buffer were added to the 450 µl sample, followed by an incubation at 37 ^0^C for 1 hour with shaking (900 RPM, Eppendorf Thermomixer), followed by adding 75 µl of 20% Triton X-100 to each tube for 1-hour incubation at 37 ^0^C with shaking (900 RPM). 20 µl samples from each tube were taken out as “undigested” and stored at -20 ^0^C. For viewpoint 1, 50 µl Csp6I (ER0211, Invitrogen) was added (500 U per tube) together with 5.6 µl 10X Buffer B (ER0211, Invitrogen) for 18 hours digestion at 37 ^0^C with shaking (700 RPM); for viewpoint 2, 10 µl DpnII (R0543M, NEB) (500 U per tube) together with 8 µl NF H2O and 2 µl 10X NEBuffer™ DpnII were added for 18 hours digestion at 37 ^0^C with shaking (700 RPM). The next day, after removing 20 µl of the sample for de-crosslinking, confirming a digestion efficiency over 80%, and performing PicoGreen DNA quantification to check the DNA concentration in each reaction, 10ug of the digested DNA was taken out into a new tube and the volume was adjusted to 600 µl with NF H2O. The samples were heat inactivated at 65 ^0^C for 20 min. Heat inactivated chromatin was added into 1X ligation buffer (EL0013, Invitrogen) supplemented with 1% Triton X-100, 0.1 mg/ml BSA, and the volume was adjusted with NF H2O to 10 ml with a final DNA concentration of 1 ng/µl. After adding 660 U T4 DNA ligase (EL0013, Invitrogen 30U/µl), samples were incubated at 16 ^0^C in thermal incubator without shaking. The next day, a final concentration of 0.5% SDS and 0.05mg/ml proteinase K (Ambion) were added to each sample, followed by 65 ^0^C incubation overnight for de-crosslinking. The next day, after adding 30 µl of RNase A (10mg/ml, PureLink), samples were incubated at 37^0^C for hour, followed by phenol: chloroform DNA purification. The chromatin was extracted with phenol:chloroform:isoamyl alcohol (25:24:1) followed by chloroform, ethanol precipitated (split to 5ml/tube and topped up with NF H2O to 15ml, then adding 100% ethanol to 68% to avoid SDS precipitation) in the presence of glycogen and dissolved in 10 mM Tris buffer (pH8). The ligated chromatin was analyzed by agarose gel electrophoresis and the concentration was determined by QUBIT HS DNA kit. 7 µg of ligated chromatin was digested with 10U specific second cutter NlaIII (R0125S, NEB) in 100 µl system with CutSmart Buffer (NEB), 37^0^C overnight without shaking. 5 µl digested chromatin was analyzed by gel electrophoresis prior to heat inactivation. Restriction enzyme was heat-inactivated by incubating the chromatin at 65°C for 20 mins. 7 µg NlaIII digested chromatin was ligated with T4 DNA ligase (EL0013, Invitrogen, 30U/µl) at 20 U/ml in 1X ligation buffer (EL0013, Invitrogen), incubated at 16°C overnight. The ligated DNA was recovered by phenol:chloroform:isoamyl alcohol (25:24:1) extraction and ethanol precipitation. 100 ng DNA of each sample was used for 4C library preparation. The library was constructed by inverse PCR and nested PCR with KAPA HiFi HotStart ReadyMix (KK2602). The 1st PCR was performed at 100 ng DNA+1.75 µl 1^st^ PCR primer mix+12.5 µl KAPA HiFi HotStart ReadyMix+H2O to 25 µl. The 1^st^ PCR program was 95°C, 3 min, 15 cycle of (98 °C, 20s; 65°C, 15s; 72 °C, 1 min), 72 °C, 5min, 4 °C hold. The 1^st^ PCR products were purified by MinElute PCR Purification Kit (28004, Qiagen) and eluted in 13 µl Elution Buffer in the kit. The 2^nd^ PCR was performed at purified 1^st^ PCR product+1.75 µl 2^nd^ PCR primer mix+12.5 µl KAPA HiFi HotStart ReadyMix+H2O to 25 µl. The 2^nd^ PCR program was 95°C, 3 min, 13 cycle of (98 °C, 20s; 65°C, 15s; 72 °C, 1 min), 72 °C, 5min, 4 °C hold. The 2^nd^ PCR products were purified by MinElute PCR Purification Kit (28004, Qiagen) and eluted in 10 µl Elution Buffer in the kit. The primer mix was 5 µl 100M forward primer + 5 µl 100M reverse primer + 90 µl H2O. All primer sequences and barcodes are listed in Supplementary Table S8. The libraries were subjected to size selection (250–600 bp) on a 4–20% TBE PAGE gel (Thermo Scientific). The TBE gel was run at 180V, 55 mins, stained with Sybr Safe and visualized with gel safe, and the libraries were extracted from PAGE using a gel crush protocol. Picogreen quantification, Bioanalyzer, and KAPA library quantification were performed to check the quality, size and amount of the recovered libraries, and NextSeq 500/550 Mid Output kit V2.5 (150 Cycles) (20024904, Illumina) was used for single end Nextseq sequencing.

### Statistical analysis

Methylation changes of clones analysed by bisulphite sequencing PCR (BSP) were calculated using the online methylation analysis tool QUMA (http://quma.cdb.riken.jp/), and the Fig. 3B was generated by R functions (http://www.r-project.org). For mRNA qRT-PCR and ChIP-qPCR, p values were calculated by t-test in GraphPad Prism Software. Values of P <0.05 were considered statistically significant (*P < 0.05; **P < 0.01; ***P < 0.001). The Mean ± SD of triplicates is reported.

### Bioinformatic analysis

#### TF bindings and motif Analysis

TF direct binding motifs surrounding p16 transcription start site were searched out using the TFregulomeR package, which is a TF motif analysis tool linking to 1,468 public TF ChIP-seq datasets in human (*33*). Specifically, we used the function *intersectPeakMatrix* from the TFregulomeR package to map the occurrences of TF motifs derived from ChIP-seq across the genomic regions of interest.

### Histone marks ChIP-Seq Analysis

Histone marks (H3K4Me3, H3K27Ac, H3K4Me1) enrichments shown in Fig. 5A were determined by ChIP-seq data cross 7 cell lines (GM12878, H1-hESC, HSMM, HUVEC, K562, NHEK, NHLF) obtained from ENCODE.

### WGBS analysis

For WGBS analysis, the leading 3 bases and adaptor sequences were trimmed from paired-end reads by *TrimGalore*. The resulting FASTQ files were analyzed by BISMARK (*34*). PCR duplicates were removed by SAMtools *rmdup* (*35*). Then *bismark_methylation_extractor* continued the extraction of the DNA methylation status on every cytosine sites. DNA methylation levels were converted into bedGraph and then to bigWig format by *bedGraphToBigWig*.

### 4C-Seq Analysis

For the 4C-seq analysis, the long-range genomic interaction regions generated by the 4C-Seq experiment were first processed using the CSI portal (*36*). Briefly, raw fq files were aligned to a masked hg19 reference (masked for the gap, repetitive and ambiguous sequences) using bwa mem (*37*). Bam files were converted to read coverage files by bedtools genomecov (*38*). The read coverage was normalized according to the sequencing depth. BedGraph files of the aligned bams were converted to bigWig format by bedGraphToBigWig. Next, the processed alignment files were analyzed using r3CSeq (*39*) and using the associated masked hg19 genome (BSgenome.Hsapiens.UCSC.hg19.masked) (*40*), from the R Bioconductor repository.

Chromosome 9 was selected as the viewpoint, and Csp6I, DpnII were used as the restriction enzyme to digest the genome. Smoothed bam coverage maps were generate using bamCoverage from the deeptools suite (*41*) with the flags “--normalizeUsing RPGC --binSize 2000 -- smoothLength 6000 --effectiveGenomeSize 2864785220 --outFileFormat bedgraph” and plotted using the Bioconductor package Sushi (*42*) to get the viewpoint coverage depth maps.

BigInteract files for UCSC and bedpe files were manually generated with the “score” values being calculated as -log (interaction_q-value_from_r3CSeq + 1*10^-10^). Sushi was then used to plot the bedpe files to get the 4C looping plots. To identify differential interaction peaks,

HOMER’s (*43*) get DifferentialPeaks was used with the flag “-F 1.5” afterwhich the corresponding bigInteract and bedpe files were generated as described.

### Data availability

All data and Supplementary Tables generated for this manuscript will be made available upon request to the corresponding authors.

## Acknowledgments

We thank Michelle Mok Meng Huang and Chelsia Qiuxia Wang for assistance with flow cytometry experiments; Melissa J. Fullwood and Ying Zhang for providing suggestions and help for the 4C experiments; S. Jha, H.P. Koeffler, and W.J. Chng for their insightful suggestions. We also thank the Tenen lab members for assistance in the experiments and reviewing the manuscript

## Funding

This research is supported by the National Research Foundation Singapore and the Singapore Ministry of Education under its Research Centres of Excellence initiative, as well as the RNA Biology Center at the Cancer Science Institute of Singapore, NUS, as part of funding under the Singapore Ministry of Education’s AcRF Tier 3 grants, Grant number MOE2014-T3-1-006. D.G.T. was funded by the Singapore Ministry of Health’s National Medical Research Council under its Singapore Translational Research (STaR) Investigator Award (MOH-STaR18nov-0002), as well as NIH grants 1R35CA197697 and P01HL131477; A.D.R. by NCI R00CA188595, the Italian Association for Cancer Research (AIRC) start up grant #2014-15347, and Fondazione Cariplo N. 2016-0476; L.C. by NIH/NHLBI grant P01HL095489; T.B. by the Canada Research Chairs program; and A.K.E. by NIH R50 CA211304

## Author contributions

D.G.T. and A.D.R. initiated the project and provided guidance throughout. Y.V.L, D.G.T, A.D.R., and T.B. designed the experiments. Y.V.L. designed and carried out experiments, analyzed data, prepared figures, and wrote the manuscript. M.K.J, Y.V.L, J.P.T, and M.T.N.L designed and carried out in vivo experiments. J.K., Q.Z. and H.K.T. carried out experiments and reviewed the manuscript. M.A.B., Q.X.X.L., and C.W. performed bioinformatics analysis and prepared figures. M.A.B., A.D.R., and L.C. edited the manuscript. T.B. and A.K.E. provided valuable suggestions about the project and reviewed the manuscript. D.G.T. and A.D.R. conceived of and supervised the project, designed experiments, and critically reviewed the manuscript.

## Competing interests

The authors declare no competing interests

## Notes

### Competing Interest Statement

The authors have declared no competing interest.

### Summary of Updates

Figure 5, along with the corresponding Result and Discussion sections, have been revised.

